# Structure-function analysis of the *Escherichia coli* β–barrel assembly enhancing protease BepA suggests a role for a self-inhibitory state

**DOI:** 10.1101/689117

**Authors:** Jack A. Bryant, Ian T. Cadby, Zhi-Soon Chong, Yana R. Sevastsyanovich, Faye C. Morris, Adam F. Cunningham, George Kritikos, Richard W. Meek, Manuel Banzhaf, Shu-Sin Chng, Ian R. Henderson, Andrew L. Lovering

**Affiliations:** Institute of Microbiology and Infection, University of Birmingham, Edgbaston, Birmingham, B15 2TT; Department of Chemistry, National University of Singapore, Singapore 117543; Institute for Molecular Bioscience, University of Queensland, St. Lucia, 4072, Australia

**Keywords:** *Escherichia coli*, structure, BepA, lipopolysaccharide, LptD, m48 metalloprotease, BAM Complex

## Abstract

The asymmetric Gram-negative outer membrane (OM) is the first line of defence for the bacteria against environmental insults and attack by antimicrobials. The key component of the OM barrier is the surface exposed lipopolysaccharide, which is transported to the surface by the essential lipopolysaccharide transport (Lpt) system. Correct folding of the Lpt system OM component, LptD, is essential and is regulated by a periplasmic metalloprotease, BepA. Here we present the crystal structure of BepA, solved to a resolution of 1.9 Å. Our structure comprises the zinc-bound m48 protease domain and a tetratricopeptide repeat (TPR) domain, consisting of four 2-helix TPR motifs and four non-TPR helices, leading to a nautilus-like shape in which the TPR repeats cup the protease domain. Using targeted mutagenesis approaches, we demonstrate that the protein is auto-regulated by the active-site plug. Further to this we reveal that mutation of a negative pocket, formed at the interface between the m48 and TPR domains, impairs BepA activity suggesting the pocket as a possible substrate binding site. We also identify a potential protein interaction site within the TPR cavity as being important for BepA function. Lastly, we provide evidence to show that increased antibiotic susceptibility in the absence of correctly functioning BepA occurs through disruption of OM lipid asymmetry, leading to reduced barrier function and increased cell permeability.

## Introduction

The outer membrane (OM) of Gram-negative bacteria is the first line of defence against environmental insults, such as antimicrobial compounds (May & Grabowicz, 2018, Nikaido, 2003). As such, the integrity of the OM must be maintained lest the bacteria become susceptible to stresses to which they would otherwise be resistant. The OM consists of an asymmetric bilayer of phospholipids and lipopolysaccharide (LPS) decorated with integral outer membrane proteins (OMPs) and peripheral lipoproteins. The impermeable nature of the OM can be attributed to several characteristics of the LPS leaflet, such as dense acyl chain packing, intermolecular bridging interactions and the presence of O-antigen carbohydrate chains (Konovalova et al., 2017, May & Grabowicz, 2018, Osborn et al., 1972, Silhavy et al., 2010, Sperandeo et al., 2017, Zgurskaya et al., 2015).

All the components required to construct the OM are synthesised in the cytoplasm. Specialised systems transport these molecules across the cell envelope and assemble the molecules into the OM in a co-ordinated fashion. Lipoproteins are transported across the periplasm and incorporated into the OM by the Lol system (Yokota et al., 1999). OMPs are transported in unfolded form across the periplasm by molecular chaperones, such as SurA and Skp, then folded and inserted into the OM by the β-barrel assembly machinery (BAM) complex. In *Escherichia coli*, the BAM complex is composed of two essential subunits, the OM β-barrel BamA and the lipoprotein BamD, and three non-essential accessory lipoproteins, BamB, BamC and BamE (Iadanza et al., 2016, Knowles et al., 2009, Mahoney et al., 2016, Wu et al., 2005). In contrast, LPS is trafficked to the OM via the Lpt system, part of which is assembled by the BAM complex (Bohl & Aihara, 2018, Chng et al., 2010a, Ma et al., 2008).

The Lpt machinery is comprised of three modules: the IM localised LptBFGC complex, which flips the LPS molecule across the IM and energises the system; LptA, which forms a bridge between the IM and OM along which the LPS travels; and the OM complex LptD/E (Bohl & Aihara, 2018, Sherman et al., 2018, Suits et al., 2008). The C-terminus of LptD forms an OM β-barrel which facilitates insertion of LPS directly into the outer leaflet of the OM (Dong et al., 2014, Qiao et al., 2014). The N-terminus encodes a periplasmic domain that interacts with the LptA bridging molecule (Chng et al., 2010b). The two LptD domains are connected via two disulphide bonds, at least one of which is required for efficient function of the LptD/E complex (Chng et al., 2012, Ruiz et al., 2010). Formation of the correct LptD disulphide bonds is reliant upon the periplasmic thiol-disulphide oxidoreductase, DsbA, as well as proper folding and insertion of the LptD β-barrel into the OM. The latter step is dependent on the BAM complex, and the interaction of LptD with its cognate OM lipoprotein partner LptE (Chng et al., 2012, Ruiz et al., 2010). To be effective at LPS delivery and to maintain the integrity of the OM, maturation of the LptD/E complex must be tightly regulated. The periplasmic chaperone DegP, the OM lipoprotein YcaL and the periplasmic metalloprotease BepA (formerly YfgC) have been implicated in controlling LptD maturation (Soltes et al., 2017).

BepA has a dual role, degrading misfolded LptD, but also promoting correct folding of the protein; loss of BepA leads to an accumulation of misfolded LptD and a reduction of mature LptD (Narita et al., 2013). Further to this, the BepA protease has been shown to interact with the main BAM complex component, BamA, which is responsible for LptD insertion into the OM. BepA can also degrade BamA under conditions of stress created by the absence of the periplasmic OMP chaperone SurA (Narita et al., 2013). BepA belongs to the m48 family of zinc metalloproteases, which are known to contain a characteristic HEXXH motif on the active-site helix (Hooper, 1994). The histidine residues within the HEXXH motif act, usually with a third amino acid and a water molecule, to coordinate the metal ion, typically zinc, at the active site (Hangauer et al., 1984, Matthews, 1988, Matthews et al., 1972). There are three other m48 family metalloprotease in *E. coli*: the OM lipoprotein LoiP, with which BepA has been shown to interact (Lutticke et al., 2012); the IM heat-shock induced endopeptidase, HtpX (Kornitzer et al., 1991); and the recently characterised OM lipoprotein YcaL, which is also involved in the regulation of LptD insertion into the OM (Soltes et al., 2017).

While there are four m48 metalloproteases in *E. coli*, BepA is the only one that has a tetratricopeptide repeat (TPR) domain at the C-terminus of the protein, with the protease domain at the N-terminus. TPR domains consist of a number of stacked repeats of α-helix pairs, together forming a solenoid-like structure that is known to facilitate protein-protein interactions and multi-protein complex formation (Perez-Riba & Itzhaki, 2019). The structure of the BepA TPR domain, from residues 310-482 of the full-length protein, was recently solved by X-ray crystallography to 1.7 Å resolution and was shown to contain 10 anti-parallel α-helices forming two subdomains each consisting of two TPR motifs. The TPR domain was demonstrated to be a site for BepA interaction with the BAM complex subunits BamA, BamD and BamC. However, the functional significance of this interaction was not investigated fully (Daimon et al., 2017).

To gain further insights into the structure-function relationship and the mechanism by which BepA acts to control and regulate folding and maturation of OMPs, particularly LptD, we sought to determine the molecular structure of the full-length protein. We present here the crystal structure of full-length BepA to a resolution of 1.9 Å, which has directed our mutagenesis approach to understand the active-site mechanism of regulation. We also present evidence of a potential substrate binding site. Finally, we demonstrate that cells lacking functional BepA exhibit disruption in OM lipid asymmetry, consistent with its crucial role in regulating LptD maturation.

## Results

### The complete BepA structure reveals a nautilus-like structure with TPR:protease contacts

We used X-ray crystallography to determine the structure of the full-length BepA protein to a resolution of 1.9 Å (data collection and refinement statistics reported in Table 1); we observe a single copy of BepA in the asymmetric unit. The structure revealed the TPR domain, consisting of 12 α-helices forming 4 TPR motifs and four non-TPR helices, in tight association with the m48 zinc-metallopeptidase domain. This forms a nautilus-like fold with the TPR subdomain cupping the metallopeptidase domain (Fig 1). During the preparation of this manuscript Shahrizal *et al*. published a crystal structure of full-length BepA to a resolution of 2.6 Å (Shahrizal *et al.*, 2019; Daimon et al 2017). These structures indicated that the BepA TPR sub-domain was composed of four TPR motif helix pairs and two non-TPR helices (nTH1 and nTH2), we have thus adopted this nomenclature. The high-resolution data presented here are in broad agreement with that presented previously, however there are some differences of note.

**Table 1.** Data collection and refinement statistics

|  | Native |
| --- | --- |
| **Data collection** |  |
| Space group | P2<sub>1</sub>2<sub>1</sub>2<sub>1</sub> |
| Cell dimensions<sup>°</sup> |  |
| *a*, *b*, *c* (Å) | 53.1, 77.0,<br>124.6 |
| Resolution (Å) | 2.18 (2.24-2.18)* |
| *R*<sub>merge</sub> | 11.1 (56.8) |
| *R*<sub>pim</sub> | 3.8 (35.8) |
| *I* / σ*I* | 20.8 (2.7) |
| CC(1/2) | 99.8 (81.7) |
| Completeness (%) | 98.9 (91.5) |
| Redundancy | 16.5 (6.0) |
| **Refinement** |  |
| Resolution (Å) | 2.18 |
| *R*<sub>work</sub> / *R*<sub>free</sub> | 20.1/24.5 |
| R.m.s. deviations |  |
| Bond lengths (Å) | 0.006 |
| Bond angles (°) | 1.00 |

\*Values in parentheses are for highest-resolution shell.

**Figure 1.**
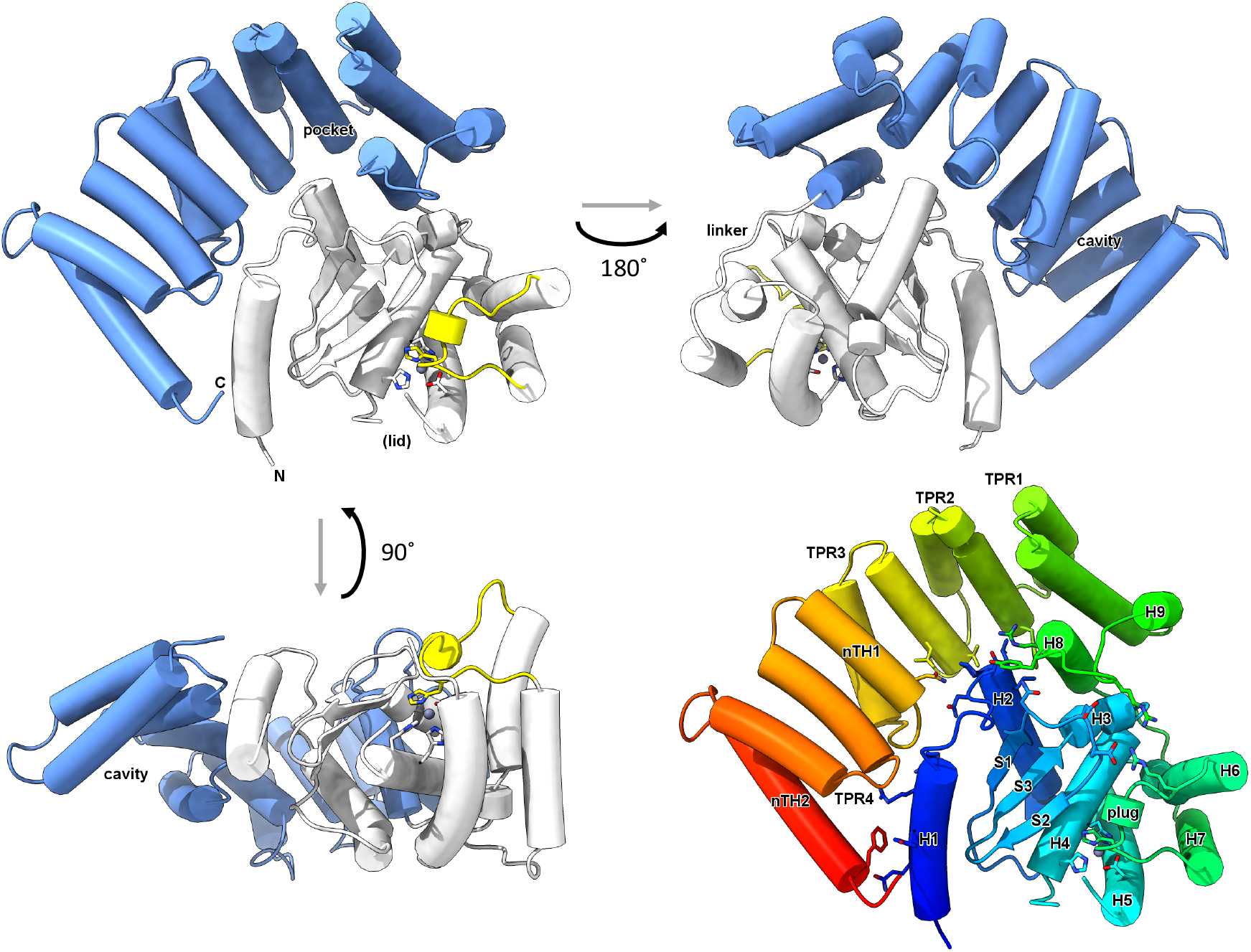
The structure of BepA reveals a nautilus-like fold. Cartoon schematic of the X-ray crystallography structure of BepA, solved to a resolution of 1.9 Å. The TPR domain is represented in blue and the protease domain in white with the active-site plug in yellow. Also labelled are the N- and C-termini, TPR pocket, TPR cavity, the linker and the site at which we expect the active-site lid (lid). Important active-site residues H136, H140, H246 and E201 are shown by stick diagram. The structure is also represented by rainbow colouring from N-to C-termini in Blue to red, respectively, with TPR motifs 1-4, non-TPR helices 1 and 1 (nTH1 and nTH2), helices, sheets and the plug labelled. Surface representation of the BepA structure coloured according to surface charge from red for negatively charged, through white for near neutral to blue for positive charge. The negatively charged TPR pocket can clearly be seen.

The structure demonstrates that the TPR domain consists of 12 α-helices, as opposed to the 10 noted previously (Daimon et al., 2017, Shahrizal et al., 2019). The reason for this difference in nomenclature is because the previously published TPR domain structure was of residues 310-482 only, comprising TPR1-nTH2, therefore excluding helices 8 and 9 (Daimon et al., 2017). We observe that helices 8 and 9 form part of the TPR domain and are preceded by an extended linker region, residues M263-S271, which connects the TPR domain to helix 7 of the protease domain (Fig 1). The additional helices of the TPR domain, helices 8 and 9, contribute a tight turn at the end of the TPR domain, allowing the m48 metallopeptidase domain to fold back and be cupped by the pocket formed by TPR motifs 2 and 3 (Fig 1). Interaction of the protease domain with the TPR pocket creates a negatively charged pocket, which is also demonstrated in the structure presented by Shahrizal *et al*. (2019). The context provided by the full-length protein structure shows that while the TPR pocket interacts with the protease domain, the TPR cavity is positioned away from the protease active-site on the opposite side (Fig 1). The cavity also comes into close proximity with the N-terminal helix, which is contained within the protease domain, therefore potentially facilitating TPR:protease domain communication.

The protease domain of BepA consists of the active-site α-helix H4 containing the HEXXH motif, and an active-site plug, which is contributed on a loop between helices H6 and H7, residues S246-P249. The TPR and protease domains are connected by the extended linker region between TPR helix H8 and protease domain helix H7, residues M263-S271, which is directly involved in positioning the active-site plug. We did not observe any density corresponding to positions L146-I194 and considering that this section is in close proximity to the active site, we expected that it may form a dynamic regulatory region (Fig 1). We sought to find evidence that the unresolved area of the protein may correspond to a dynamic lid. Therefore, we scrutinised the Protein Data Bank (PDB) for similar structures. Information on the missing region of our structure can be inferred from an unpublished structure in the PDB of an m48 zinc-metallopeptidase from *Geobacter sulfurreducens*, which consists of only the protease domain, with no associated TPR (PDB: 3C37). The structure of the *Geobacter* protease structure provides some information on the missing section and demonstrates a short three-turn extension to the C-terminus of active site helix H4, beyond that seen in the BepA structure. This is followed by a glycine facilitated kink and another three helical turns terminating at residue D136 of the 3C37 structure (Appendix Fig S1). The *Geobacter* is also missing a section, D136-N139, however residues M140-F149 form another short α-helical region, which is connected to the N-terminus of helix H5, by an extended region formed by residues G150-S158 of the 3C37 structure (Appendix Fig S1). The recently published BepA structure also provides some information on the BepA active-site lid, demonstrating a similar extension of active-site helix H4, seen in PDB: 3C37, and a helix from residues T160-Q176, which is on a slightly different angle to that of 3C37, residues M140-F149 (Shahrizal et al., 2019). Overall, comparison of the structure presented here, that of Shahrizal *et al*. (2019) (PDB: 6AIT), and the *G. sulfurreducens* structure (PDB: 3C37), suggests that the missing section from the structure presented here may form a putative active-site lid. The putative lid, along with the plug, likely regulates access to the active-site. The fact that no density for the putative lid is observed in our structure, and that sections are missing in those presented previously, suggests that the active-site lid is dynamic and may adopt multiple conformations.

### Functional analysis of the BepA active-site suggests an auto-inhibitory state

The structure shows the HEXXH motif, which is characteristic of zinc-dependent metallopeptidases (Hooper, 1994, Matthews, 1988) and, in this case, it is found within helix H4 (Fig 1 and Fig 2A). The active-site zinc ion is coordinated by H136 and H140 within the HEXXH motif, E201 contributed by helix H5, and H246 on a loop that forms the small α-helical active-site plug (Fig 1 and Fig 2A). This is informative as in general Zn-dependent proteases, the active-site zinc ion is usually chelated by three amino acid residues and one water molecule, which is normally utilised to catalyse proteolysis of the substrate (Hangauer et al., 1984, Matthews et al., 1972). Co-ordination of the zinc ion in this manner fulfils the fourth ligand, which is an amino acid instead of a water molecule and is reminiscent of an auto-inhibitory metamorphic zinc metallopeptidase encoded by *Methanocaldococcus jannaschii* (Lopez-Pelegrin et al., 2014). This zinc metallopeptidase exists as a proteolytically active monomer at lower concentrations, but transitions to an inactive dimer and tetramer at higher concentrations, in which the third zinc-coordinating histidine is contributed by the dimer interaction in a conformation that resembles that of the BepA active-site plug seen in our crystal structure (Fig 2). The active-site plug also contributes hydrophobic residues I242 and L243, which likely fulfil interactions with other hydrophobic residues in the active-site, such as V133, F225 and L229. Further to this, alignment with the human nuclear membrane zinc metalloprotease ZMPSTE24 structure with a trapped substrate (PDB: 2YPT) reveals that the BepA active-site plug occupies the same physical space as the substrate for ZMPSTE24 (Fig 2B). Residue H246 on the BepA active-site plug directly clashes with positioning of substrate in the 2YPT structure and the hydrophobic residues I242 and L243 occupy a similar space to the 2YPT substrate hydrophobic residues I3’ and M4’ (Fig 2B). Based on these observations, we anticipated that H246, which is contributed by the active-site plug, is likely to relocate to unplug the active-site in order to facilitate protease activity of BepA at least to a position similar to H459’ in the 2YPT structure (Fig 2B). Therefore, we hypothesised that the active-site plug sterically hinders interaction of the substrate with the BepA active-site, effectively auto-inhibiting BepA proteolytic activity. Also, the active-site plug features directly after the TPR domain in the linear structure and is therefore connected in such a way to move in response to a signal generated through substrate interaction, potentially sensed by the TPR (Fig 1 and Fig 2A).

**Figure 2.**
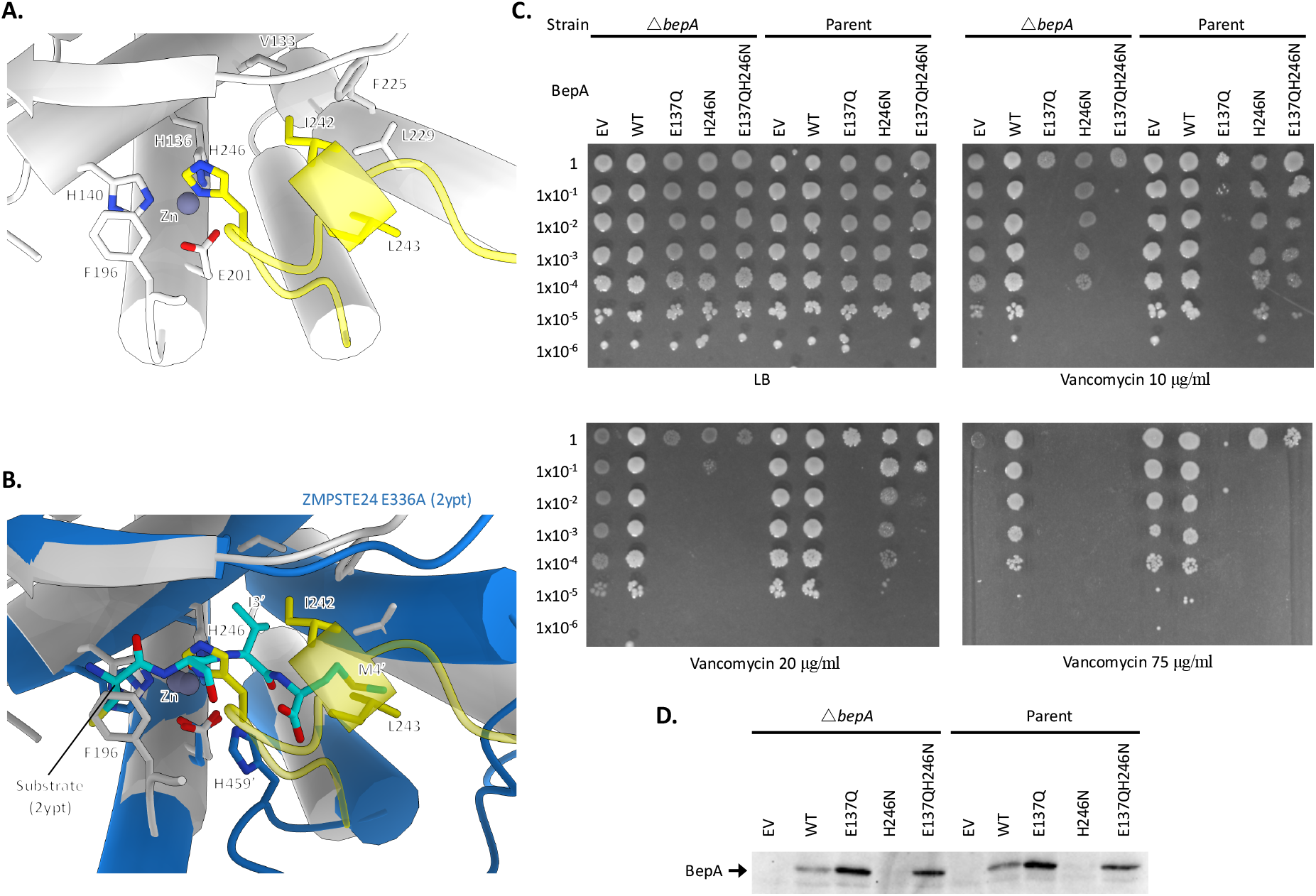
The BepA active-site plug acts to regulate BepA proteolytic activity. Analysis of the BepA structure suggested a regulatory role for the active-site plug, therefore plasmids encoding mutated *bepA* were screened for their capacity to complement the vancomycin sensitivity of ΔbepA E. coli. **A.** Structural diagram of the BepA active-site with key residues represented by stick diagram and labelling. **B.** Alignment of the BepA active-site (transparent white and yellow ribbon) with that of the human nuclear membrane zinc metalloprotease ZMPSTE24 mutant E336A with a synthetic substrate peptide (PDB: 2YPT) (Quigley et al., 2013) (Opaque blue ribbon) 2YPT residues are labelled with the addition of a ‘ symbol. **C.** Screen for vancomycin sensitivity of cells carrying pET20b encoding WT or mutated copies of BepA in the parent or Δ*bepA* strain background. The empty vector control is labelled EV. Cells are normalised to OD_600_ = 1 and ten-fold serially diluted before being spotted on the LB agar containing the indicated antibiotics (all plates contain 100 μg/ml carbenicillin additionally). **D.** Western blot using anti-6xHis antibody raised in mice and anti-mouse::HRP secondary antibody to target the BepA::6xHis protein in samples used for vancomycin sensitivity screens.

To test the importance of H246 in occupying the fourth coordination site on the zinc ion, we generated a conservative mutation of the H246 position to asparagine (H246N). The aim was to conserve structure of the active-site plug, but abolish coordination of the zinc ion by this residue in order to “de-regulate” the protease activity (Lopez-Pelegrin et al., 2014). We also constructed the E137Q mutation in the active helix HEXXH motif, which is known to abolish protease activity of BepA and would allow us to compare the activities of the mutated proteins (Narita et al., 2013). To test whether the H246N BepA mutant is functional, we assayed the ability of this mutant to complement the Δ*bepA* strain, which is known to exhibit increased sensitivity to large antibiotics such as vancomycin, presumably due to impaired barrier function of the OM. The E137Q active-site mutant was incapable of restoring vancomycin resistance to Δ*bepA* cells and had a severe negative effect on the growth of the Δ*bepA* mutant (Fig 2C). Furthermore, we observed that the H246N mutant BepA was also incapable of complementing vancomycin sensitivity of the Δ*bepA* cells; however, while the H246N protein also severely increased the vancomycin sensitivity of the mutant, the negative effect was less extreme than with the E137Q version of the protein (Fig 2). Considering this phenotype, we decided to investigate if the mutated proteins had a dominant-negative effect in the parent background expressing wild-type *bepA*. We found that the empty vector and wild-type BepA had no detrimental effect on BW25113 parent cells grown in the presence of vancomycin. Our analysis of the E137Q mutant was in agreement with previous studies when analysed in the parent background and demonstrated a severe dominant-negative phenotype (Narita et al., 2013). We also observed that the presence of H246N BepA had a dominant-negative effect on the capacity of the cells to grow in the presence of vancomycin, despite the presence of wild-type BepA expressed from the chromosomal locus. Similarly to the effect in the mutant background, the dominant-negative effect of the H246N protein was less severe than that of the E137Q mutant. We speculate that this is because the active-site plug is less able to interact with the active-site zinc ion, as observed in our crystal structure of BepA, and that the protein may be in a constitutively activated, or “de-regulated”, conformation (Fig 1 and Fig 2). We used western blotting to detect the expression of BepA proteins in whole cell lysates using anti-6xHis antibodies. The western blot showed an elevated level of the E137Q protein compared to wild-type and an absence of observable tagged protein in the H246N sample (Fig 2D). These observations were consistent between the Δ*bepA* and parent backgrounds (Fig 2D and Appendix Fig S2). These results support the hypothesis that the E137Q mutation renders BepA protease inactive, therefore stabilising the protein due to a lack of auto-proteolytic activity. Considering that we know the H246N BepA is expressed, as it has a dominant-negative effect, the absence of a detectable tagged protein by western blot suggests the His-tag may be degraded, supporting the hypothesis that this mutation gives rise to a protein with de-regulated proteolytic activity.

In order to test if the dominant-negative effects of the H246N protein were due to increased protease activity, we tested whether the established protease dead mutation, E137Q, could alleviate the dominant-negative effect of the H246N mutation or if the negative effects would be synergistic. As expected, BepA E137Q H246N was not only incapable of complementing the vancomycin sensitivity, but had a more severe dominant-negative effect than the H246N mutation alone, similar to that of E137Q (Fig 2C). The dominant-negative effect of the double mutant is likely due to the E137Q mutation. We also analysed expression of the E137Q H246N BepA protein by western blot and found a similar level of tagged protein compared to the E137Q protein (Fig 2D). This suggests that the potential auto-proteolytic activity created by the H246N mutation was alleviated by the introduction of the E137Q protease-dead mutation. These results also imply that the dominant-negative effect of the E137Q and E137Q H246N mutations is unlikely due to un-regulated protease activity and more likely due to inappropriate interactions with substrate, or the chaperone activity of BepA.

### Mobility of the active-site plug is required for BepA function

We hypothesised that the conformation of BepA observed in our crystal structure is in the inactive form and that movement of the active-site loop “unplugs” the active-site in response to some unknown signal. To test this hypothesis, we aimed to “lock” the active-site in an inhibited state by engineering a disulphide bond. Cysteine substitutions were introduced into proximal sites in BepA, specifically at positions E103, in the loop between S1 and S2, and E241 in the active-site plug, either individually or in concert (Fig 3A and Fig 3B). The single cysteine substitutions complemented the sensitivity phenotype, indicating that the single substitutions had no impact on BepA function. However, the double cysteine mutant was incapable of restoring vancomycin resistance to the *bepA* mutant under normal growth conditions. In contrast, in the presence of the reducing agent TCEP (tris(2-carboxyethyl)phosphine), the double cysteine mutant was able to complement vancomycin sensitivity (Fig 3C). These observations suggest that in the double-cysteine-containing BepA a disulphide bond was formed that “locked” the active-site loop in the inactive conformation preventing BepA from promoting LptD maturation. This inhibition was alleviated by breaking the disulphide bond in the presence of TCEP, therefore allowing free movement of the regulatory active-site plug and normal functioning of BepA (Fig 3C). As would be expected of an inactive “locked” BepA mutant, the double-cysteine mutant was much more sensitive to vancomycin only in the absence of TCEP, whether in a *bepA* or wild-type background, indicative of dominant-negative effects similar to E137Q mutation (Fig 3C).

**Figure 3.**
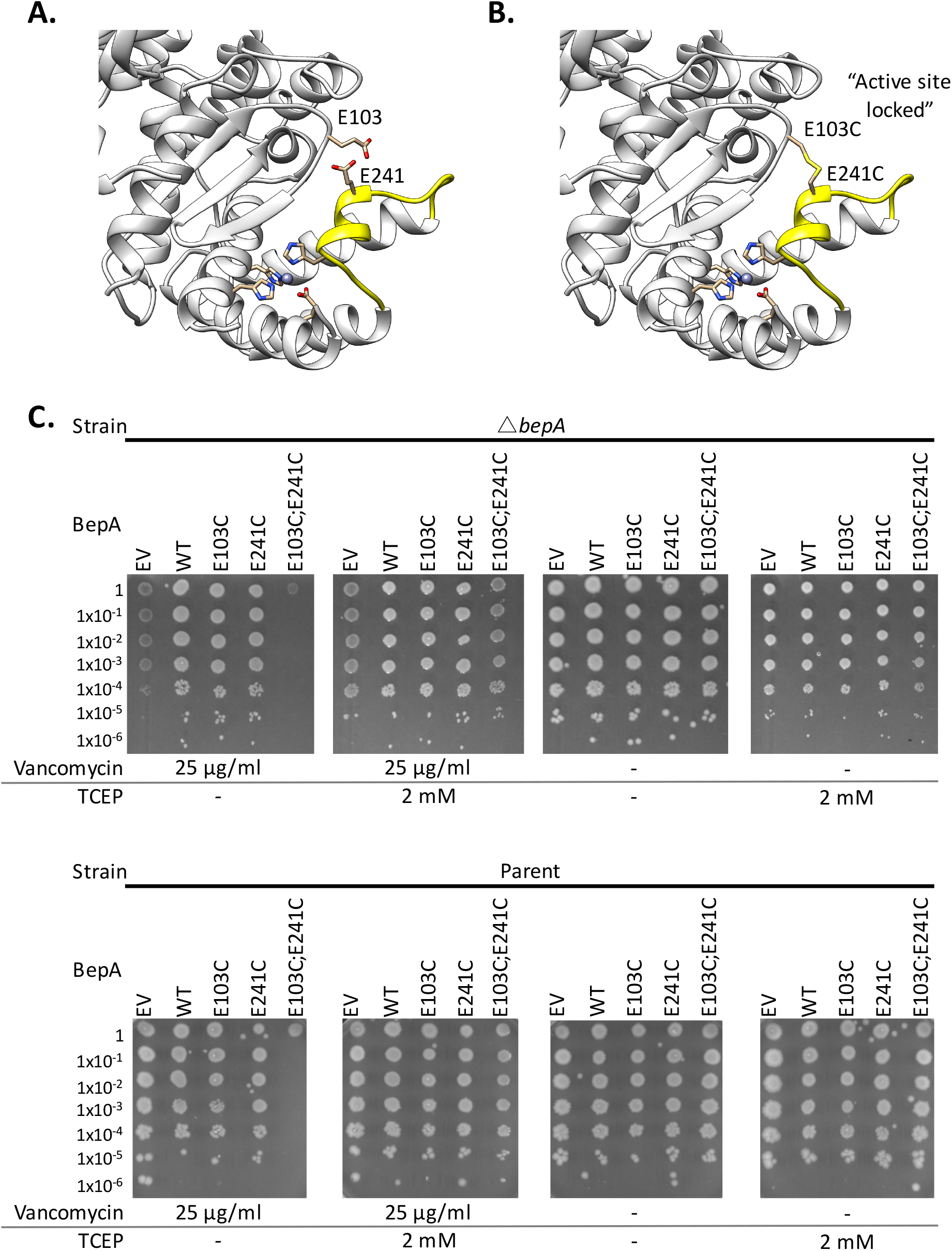
Flexibility of the active-site plug is required for BepA function. The requirement for flexibility of the BepA active-site plug for full BepA function was assayed by disulphide bond locking of the active-site plug and functional screening. **A.** Structural representation of the BepA active-site with residues targeted for mutation to cysteine, E103 and E241, labelled and coloured yellow. **B.** Structural representation as in A. except with the E103 and E241 residues mutated to cysteine and bonded in what is expected to be the “active-site locked” conformation. **C.** Screen for BepA function and disulphide bond “locking” of the BepA active-site plug to the main protease domain completed vancomycin sensitivity screening with the addition of the reducing agent TCEP where indicated.

### BepA sub-domains communicate and interact with substrate via a negatively charged cleft

The TPR domain contains two potential substrate binding sites, termed the “pocket” on the large palm and the “cavity” on the small palm (Fig 1 and Fig 4). We identified two conserved charged residues, R280 and D347, in the BepA TPR pocket, which in the complete structure forms a larger negatively charged cleft through interaction with the protease domain (Fig 1 and Fig 4A). This negatively charged cleft is connected to the active site via a negatively charged ditch and so has been hypothesised to facilitate substrate interactions; however, no evidence for substrate interaction or importance for BepA function has yet been provided (Fig 4A). The R280 and D347 residues form a salt bridge that appears to stabilize interactions between the TPR and protease domains (Fig. 4A). We targeted these two conserved charged residues within the pocket, and sought to assess their requirement for BepA function by mutational analysis and functional screening. Our analyses revealed that the R280 mutations had no appreciable effect on BepA activity. However, the D347R mutation had a negative effect on the capacity of the BepA protein to complement the vancomycin sensitivity of the *bepA* mutant, despite the proteins being expressed (Fig 4B and Appendix Fig S2). We also found that the D347R mutation had a dominant-negative effect in the parent background (Fig 4B). The D347R mutation likely disrupts the negatively charged pocket, therefore preventing the negatively charged pocket from directing the substrate to the prominent positively charged R280 within the pocket (Fig 4A).

**Figure 4.**
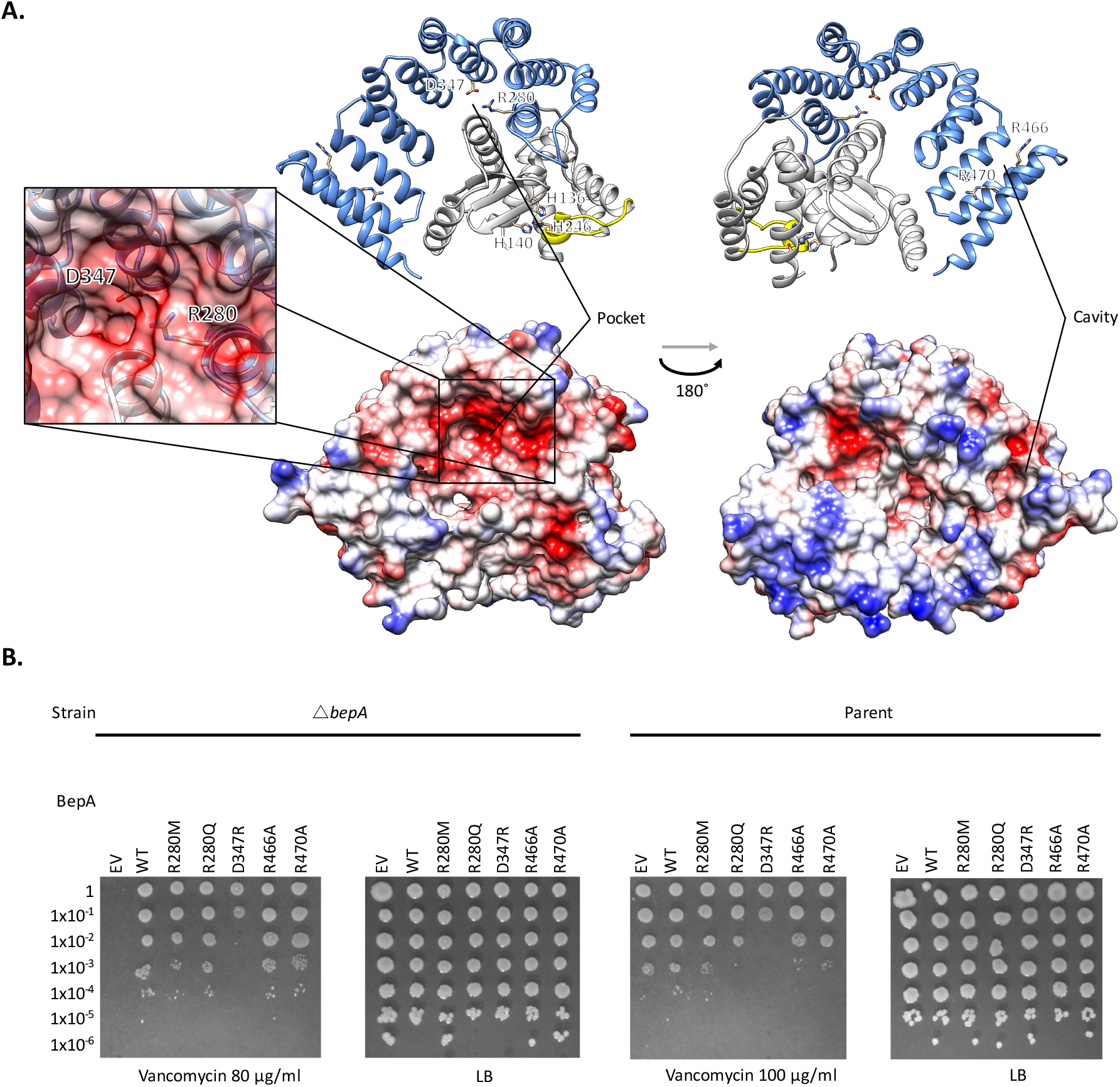
The BepA negatively charged pocket is a potential substrate binding site. Interaction of the TPR domain with the protease domain of BepA creates a negatively charged pocket containing evolutionary conserved residues, which were mutated to test their importance for BepA function. **A**. Ribbon diagram of the BepA crystal structure with key evolutionarily conserved residues represented as stick diagrams and labelled. Also shown are surface representations of BepA in the same orientations coloured according to surface charge from red for negatively charged, through white for near neutral to blue for positive charge. The negatively charged TPR pocket can clearly be seen. The zoom-in box shows the negatively charged pocket with the surface made transparent to demonstrate the position of the key D347 and R280 residues. **B**. Screen for vancomycin sensitivity of cells carrying pET20b encoding WT or mutant copies of BepA in the parent or Δ*bepA* strain background. The empty vector control is labelled EV. Cells are normalised to OD_600_ = 1 and ten-fold serially diluted before being spotted on the LB agar containing the indicated antibiotics (all plates contain 100 μg/ml carbenicilin additionally).

We next sought to assess the cavity in the TPR domain, in which conservation analysis revealed two conserved arginine residues R466 and R470. We expected these residues might be involved in substrate recognition or interaction with protein complex partners due to the prominent position in the cavity and their high level of conservation despite any obvious structural role (Fig 4A). Therefore, we mutated these residues to alanine in order to assess their contribution to the function of BepA. Unexpectedly, mutation of these residues appeared to have no impact on the function of the BepA protein (Fig 4B). We expect that the cavity of BepA might still play a role in protein interaction and substrate recognition, although more extensive mutagenesis will be required to further elucidate this function while using the vancomycin sensitivity screen.

### Loss of BepA leads to increased OM permeability

It has been established that *bepA* mutant *E. coli* are more sensitive to antibiotics with a high molecular mass, such as vancomycin, erythromycin, rifampicin and novobiocin (Narita et al., 2013). This is presumed to be due to increased OM permeability. The hypothesis is that the loss of BepA results in reduced LptD assembly, therefore leading to reduced OM LPS content. This would in turn cause phospholipids to flip from the inner leaflet to the outer leaflet of the OM, creating a perturbation in OM lipid asymmetry and increased OM permeability to large antibiotics (Narita et al., 2013, Ruiz et al., 2009, Soltes et al., 2017). However, antibiotic sensitivities alone do not necessarily support this hypothesis directly and therefore we sought to test this hypothesis experimentally.

In order to test the permeability of the *bepA* mutant compared with parent BW25113 cells, we used the chlorophenyl red-β-D-galactopyranoside (CPRG) assay. CPRG is a β-galactosidase substrate, however it fails to penetrate wild-type *E. coli*, therefore being inaccessible to cytoplasmic β-galactosidase, which would normally hydrolyse the CPRG and produce a red colour (Kritikos et al., 2017, Paradis-Bleau et al., 2014). Production of the red colour is an indicator of cell permeability and thus can be measured using a time resolved wavescan of cells grown on LB agar supplemented with CPRG. The BW25113 parent strain is Lac^−^, therefore the strains were co-transformed with the relevant *bepA* encoding plasmid and a *lacZYA* expression vector, pRW50 CC-61.5 (Gaston et al., 1990, Kritikos et al., 2017, Paradis-Bleau et al., 2014). CPRG assays indicated that the *bepA* mutant cells are indeed more permeable to the β-galactosidase substrate CPRG and that this permeability phenotype can be complemented with leaky expression of *bepA* at low levels from the pET22b plasmid (Fig 5B). Active site mutants that result in inactive (E137Q) or presumably de-regulated (H248N) BepA are not able to restore the OM barrier against CPRG. In addition, mutations altering conserved residues in either the pocket (R280, D347R) or cavity (R460, R470) are also not able to fully complement the permeability defect. This is particularly surprising since the R280, R460 and R470 mutations have very strong negative effects on cell permeability, despite there being no observed effect of these mutations in the vancomycin sensitivity assay (Fig 4B and Fig 5). This may suggest that the R460 and R470 mutations affect BepA function in a way that leads to CPRG specific increased permeability or alternatively that the phenotypes caused by these mutations are much milder than the active site mutations. The mild permeability phenotype could explain the lack of vancomycin sensitivity but with increased permeability to the much smaller CPRG dye. These results also imply that the BepA TPR cavity could be involved in protein interactions or even be a substrate binding site.

**Figure 5.**
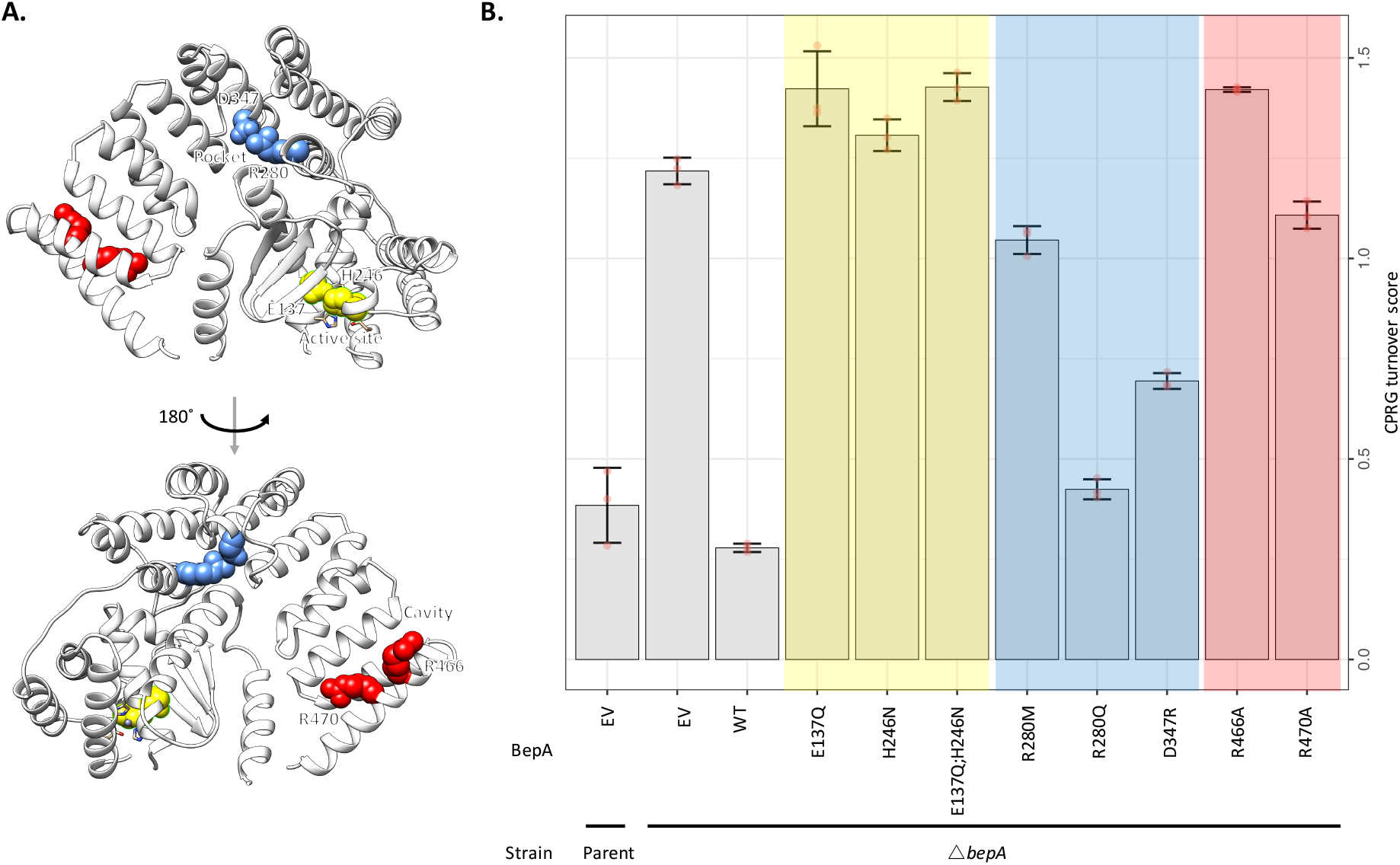
Loss of BepA leads to increased cell permeability. Cells in which *bepA* had been deleted were complemented with plasmids encoding mutated copies of *bepA* and subjected to permeability tests using the β-galactosidase substrate Chlorophenol red-β-D-galactopyranoside (CPRG), to which a wild-type *E. coli* OM is not normally permeable. **A.** Structure of BepA showing residues targeted for mutation as colour-coded spheres that match the colour-coded chart represented in panel B. **B.** CPRG permeability assay of Parent or Δ*bepA* cells carrying pET20b with WT or mutant copies of BepA as indicated. The empty vector control is labelled as EV. Change in absorbance at OD_575_ compared to the Lac^−^ cells is represented for two independent experiments each containing three replicates.

### Increased permeability is due to increased surface exposed phospholipid

While we have established that the Δ*bepA* cells are more permeable, this is not necessarily evidence of perturbed OM lipid asymmetry. Perturbation of OM lipid asymmetry can be detected through monitoring the activity of the enzyme PagP. On detecting surface exposed phospholipids, the OM localised Lipid A palmitoyltransferase PagP, utilises the outer leaflet phosphoplipids as palmitate donors to convert hexa-acylated Lipid A to hepta-acylated Lipid A (Bishop et al., 2000, Dekker, 2000, Hwang et al., 2002). The resulting lyso-phospholipid product is then degraded by the OM phospholipase PldA (Fig 6A). To measure the levels of hepta-acylated Lipid A, radiolabelled Lipid A was isolated from the Δ*bepA* mutant or bacteria that had been complemented with BepA, BepA E137Q or BepA H246N. The lipids were then separated by thin layer chromatography. The parent strain BW25113, transformed with empty pET22b, were treated with EDTA prior to Lipid A isolation, a process that is known to induce high levels of hepta-acylated Lipid A production and act as a positive control (Chong et al., 2015, Jia et al., 2004, Yeow et al., 2018). As expected, Δ*bepA* mutant cells showed a significant increase in the levels of hepta-acylated Lipid A in relation to hexa-acylated Lipid A, indicating perturbation of OM lipid asymmetry in the absence of functional BepA (Fig 6). While the catalytically dead E137Q and the “de-regulated” H246N mutants were not able to rescue this defect, they also did not appear to significantly increase the levels of hepta-acylated Lipid A compared to cells lacking BepA; this is surprising given that both these mutations cause severely increased vancomycin sensitivity (Fig 6). Additionally, we did not see any effect on lipid A palmitoylation for any of the other mutations used in this study. Essentially, there seems to be very little correlation between the extent of OM lipid asymmetry defect and vancomycin sensitivity/CPRG permeability. Together, our results demonstrate that cells exhibit defects in OM lipid asymmetry in the absence of BepA, which only contribute in part to some of the permeability and/or antibiotic sensitivity defects in these strains.

**Figure 6.**
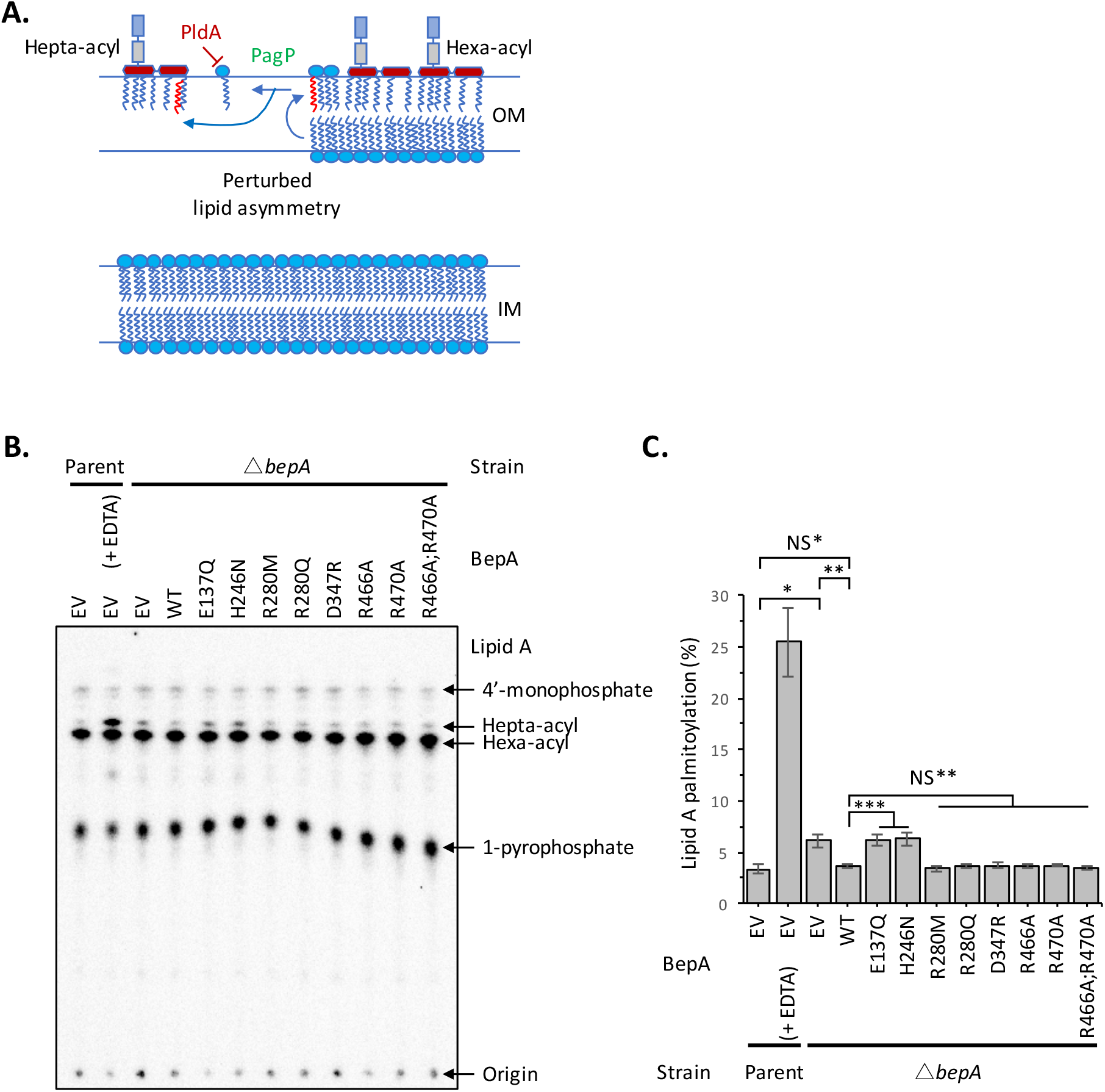
BepA mutants are more permeable due to surface exposed phospholipid. The increased permeability of ΔbepA cells was hypothesised to be due to increased surface exposed phospholipid, therefore this was tested by the PagP mediated Lipid A palmitoylation assay, which detects surface exposed phospholipid. **A.** Schematic demonstrating the role of PagP in sensing and responding to surface exposed phospholipid. **B.** PagP mediated Lipid A palmitoylation assay. PagP transfers an acyl chain from surface exposed phospholipid to hexa-acylated Lipid A to form hepta-acylated Lipid A. [32-P]-labelled Lipid A was purified from cells grown to mid-exponential phase in LB broth with aeration. Equal amounts of radioactive material (cpm/lane) was loaded on each spot and separated by thin-layer chromatography before quantification. As a positive control, cells were exposed to 25 mM EDTA for 10 min prior to Lipid A extraction in order to chelate Mg^2+^ ions and destabilise the LPS layer, leading to high levels of Lipid A palmitoylation. **C.** Hepta-acylated and hexa-acylated lipid A was quantified and hepta-acylated Lipid A represented as a percentage of total. Triplicate experiments were utilised to calculate averages and standard deviations with students t-tests used to assess significance. Student’s *t*-tests: \**P* < 0.005 significant compared with Parent EV; \*\**P* < 0.005 significant compared with ΔbepA EV; *** *P* < 0.001 significant compared with ΔbepA WT; NS* *P* > 0.1 compared with Parent EV; NS** *P* > 0.1 compared with ΔbepA WT.

## Discussion

In this study we present the structure of full-length BepA at a resolution of 1.9 Å, which is a periplasmic m48 zinc metalloprotease family protein involved in regulating the maturation of the LPS biogenesis machinery in Gram-negative bacteria. The structure has guided our mutagenesis strategy to investigate the auto-regulatory nature of the BepA active-site and has identified a potential substrate interaction site and a protein interaction site. Loss of the BepA protein is known to cause *E. coli* cells to become hypersensitive to antibiotics and antimicrobial products that have a large molecular mass, or are particularly hydrophobic, and this is hypothesised to be due to increased cell permeability (Daimon et al., 2017, Narita et al., 2013, Shahrizal et al., 2019, Soltes et al., 2017). Here we have shown by the use of a CPRG permeability assay that lack of BepA causes *E. coli* cells to become more permeable due to a reduction in the efficacy of the OM permeability barrier, which is in part due to an increase in the quantity of surface exposed phospholipid (Malinverni & Silhavy, 2009, Nikaido, 2005). The OM of *E. coli* is asymmetric with phospholipid forming the inner leaflet and LPS forming the outer leaflet. Presumably, loss of BepA leads to a decrease in productive LPS biogenesis machinery and therefore a decrease in OM LPS content, leading to phospholipid migrating from the inner to the outer leaflet of the OM in order to balance the membrane. Further support of this hypothesis comes from the supplementary dataset presented by Narita *et al.* (2013), in which they sought multi-copy suppressors of the antibiotic sensitivity created by loss of BepA. Interestingly, the enzyme PagP, which is the subject of the PagP assay used here, was found as a multi-copy suppressor of the Δ*bepA* mutant hypersensitivity to erythromycin and vancomycin. PagP is responsible for reacting to surface exposed phospholipid by using it as a palmitate donor for acylation of Lipid A, therefore increasing the expression of PagP in the absence of BepA might help to alleviate the problem of surface exposed phospholipid.

The data presented here suggest that the defects in lipid asymmetry are not wholly responsible for the observed increases in vancomycin sensitivity. This is not entirely unexpected as certain mutants in the maintenance of lipid asymmetry (Mla) pathway actually demonstrate increased vancomycin sensitivity despite an accumulation of surface exposed phospholipid (Isom et al., 2017). Therefore, there may be other reasons for the increase in vancomycin sensitivity. One potential explanation being that in the absence of BepA, or presence of mutated BepA, LptD barrels are more often stalled in a rudimentary form that could weaken the OM permeability barrier to vancomycin. The work of Soltes *et al.* (2017) suggests that BepA specifically targets partially folded rudimentary LptD β-barrels that are still associated with the BAM complex, as opposed to periplasmic substrates that have fallen off the pathway or very early BAM-associated unfolded substrates, which are targeted by DegP and YcaL respectively. In order to achieve this, Soltes *et al*. showed that BepA specifically targets the *LptD4213* mutant, which has a 23 amino acid deletion in LptD and causing LptD to become stalled while partially folded around LptE and remains in contact with BamA and BamD (Lee et al., 2018, Lee et al., 2016). This work provides evidence to support the idea that non-functional BepA could lead to an accumulation of partially folded LptD barrels, which could in turn facilitate the entrance of vancomycin into the cell. These results and the mutants presented here might be informative in trying to understand the order in which BepA interacts with and recognises the BAM complex and misfolded LptD substrates. Considering that this is a confirmed stalled target of BepA, the “protease dead” mutant reported here may act to further enhance the detrimental effects caused by this stalled mutant. Further studies utilising the misfolded intermediates at the disposal of Soltes *et al*., may yield further clarity on this subject.

The structure presented here allowed us to make assumptions about how the active-site of BepA might be regulated. Comparison of BepA with the structure of human nuclear membrane zinc metalloprotease ZMPSTE24 containing a trapped substrate (PDB: 2YPT) demonstrated that the BepA active-site plug occupies the space that should be occupied by substrate. This therefore implied that the active-site plug serves an auto-inhibitory role and would likely have to relocate in order to facilitate BepA activity. It therefore stood to reason that the active-site plug should be flexible, thus we sought to disrupt the interaction between the active-site plug and the chelated zinc ion. We found that weakening the interaction of the active-site plug with the zinc ion led to a significant dominant-negative effect on wild-type cell vancomycin sensitivity, presumably due to the protein becoming “de-regulated”. This is likely because of the lack of auto-inhibition, as seen with the inactive dimer of the *M. janaschii* protein, which has a very similar arrangement around the active-site zinc (Lopez-Pelegrin et al., 2014). Interestingly, we found that both weakening this interaction, and forcing the active-site plug to be locked in place by introduction of cysteine mutations and putative disulphide bonding, caused a dominant-negative effect on parent cells. This not only confirms that the active-site plug is flexible and must relocate in order for substrate to be accommodated in the active-site, as hypothesised previously (Shahrizal et al., 2019), but also that if the active-site plug is locked in place then the BepA protein interacts with substrate or partner proteins in a way that is detrimental to the cells. This effect is similar to, but not as extreme as, the severe dominant-negative effect of the “protease-dead” mutation E137Q, presented previously (Narita et al., 2013). We have provided evidence that this mutation prevents auto-proteolytic activity of BepA, therefore supporting the idea that this mutation causes the loss of BepA proteolytic activity. We hypothesise that these mutations may have an effect on the chaperone activity of BepA, or may lock the protein into an inactive complex with the BAM machinery. In this scenario, the “de-regulated” activity of the H246N BepA would allow it to escape such a complex and even complete the appropriate proteolytic activity on occasion, whereas the “protease-dead” mutations could lead to the protein becoming trapped.

During the preparation of this manuscript, a structure of the BepA protein was released in which the negatively charged pocket formed between the TPR and protease domains was suggested as the likely candidate for substrate recognition, however no evidence for substrate binding or an effect on BepA function was provided (Shahrizal et al., 2019). Evolutionary conservation analysis directed us to this site independently, as being important for BepA function. Our analysis of the BepA structure presented here identified residues D347 and R280 within the negatively charged pocket as key residues. Our targeted mutagenesis confirmed that both of these residues are important for the correct function of BepA, therefore providing further support for this as the potential site of substrate interaction. Our conservation analysis also identified two conserved arginines R460 and R470 in the TPR cavity as potentially important for function. Interestingly, we found no effect of mutating these residues when BepA function was assayed by vancomycin sensitivity screen, however the CPRG permeability assay suggested that these residues were particularly important. The TPR cavity has previously been implicated as a protein binding site through photo-crosslinking experiments, that show the BepA TPR cavity interacts with BamA, BamD and BamC (Daimon et al., 2017). This function is in agreement with the known role of TPR domains in numerous other proteins in which they serve as interaction domains to facilitate the formation of protein complexes (Zeytuni & Zarivach, 2012). Taken together, these results suggest the TPR cavity as a second potential protein binding site and one can foresee a situation where BepA needs to interact with both the BAM machinery and recognise misfolded/stalled rudimentary LptD β-barrels in order to function correctly, with the TPR cavity acting as a BAM complex docking site and the pocket as a substrate recognition site. This hypothesis is in agreement with the suggestion that BepA reversibly interacts with the BAM complex and the partially folded substrate, depending on the identity and folding state of the substrate (Soltes et al., 2017). Alternatively, the TPR cavity may play a role in the proposed chaperone activity of BepA (Narita et al., 2013).

Our work utilised a high-resolution structure of BepA to direct a study to understand the mechanistic function of the BepA protease active-site and to identify two potential protein binding sites. Further to this, we demonstrate that the function of the BepA active-site is sensitive to perturbation in two separate ways, either by increasing flexibility of the regulatory active-site loop or by restricting the loops motion. These two perturbations lead to severe permeability defects beyond that achieved by the loss of BepA and causes the cells to become sensitive to antibiotics to which Gram-negative bacteria are normally resistant. Therefore, this highlights BepA as a potential future drug target which would allow combination therapies to increase the efficacy of, or enable the use of, certain antibiotics to which Gram-negative bacteria are currently resistant.

## Methods

### Expression and purification of BepA

The BepA open reading frame, including N-terminal signal peptide, was codon optimised for expression in *E. coli* and cloned into the IPTG inducible vector pET20b fused to a C-terminal His_6_-tag (a service provided by Genscript). This vector was transformed into *E. coli* DE3 cells and used for recombinant protein production. Briefly, overnight cultures grown in LB media at 37°C were used as the inoculum for auto-induction media supplemented with 10 µM ZnCl_2_. The resulting cultures were grown at 37°C to an OD_600_ of ~0.8 before the temperature was changed to 18°C for ~18 hours. Cells were harvested by centrifugation and cell pellets were stored at −80°C.

To purify His-tagged BepA, cell pellets were resuspended in buffer A (20 mM imidazole, pH 7.5; 400 mM NaCl) supplemented with 0.05% Tween20 and lysed by sonication. Cell lysates were clarified by ultra-centrifugation and then incubated with Super Ni-NTA agarose resin (Generon) at 4°C with gentle agitation overnight. The incubation mixture was centrifuged briefly, the supernatant was removed, and the resin was resuspended in buffer A before being loaded onto a gravity-flow purification column. The resin was washed extensively with buffer A, then with 20 ml of Buffer A supplemented with 50 mM imidazole, before washing with buffer B (400 mM imidazole, pH 7.5; 400 mM NaCl; 2 % glycerol). BepA protein, eluted in buffer B, was dialysed against buffer C (20 mM MES, pH 6.5; 5 mM EDTA) at 18°C for 6 h (to remove metals co-purified with BepA protein) and then dialysed, extensively with sequential buffer changes, against buffer D (as buffer C but lacking EDTA and instead supplemented with 10 µM ZnCl_2_ and 150 mM LiSO_4_) at 18°C. BepA protein was concentrated to ~60 mg/ml by ultra-filtration and then further purified on a HiLoad Superdex 200 26/600 column (GE Healthcare) equilibrated in buffer D. Fractions containing pure BepA protein were pooled and concentrated to 35 mg/ml for use in crystallization trials.

### Crystallisation and determination of BepA structure

Purified recombinant BepA was used with proprietary crystal screens (supplied by Molecular Dimensions and Jena Bioscience) in sitting drop crystallization experiments using 2 µl of protein solution and 2 µl of crystallization mother liquor at 18°C. Large crystals were obtained in 0.1 M Na HEPES, pH 7.0, and 8% w/v PEG 8,000 and grew within 30 days. Crystals were cryo-protected by step-wise addition of mother liquor supplemented with 25 % ethylene glycol prior to flash freezing in liquid nitrogen.

Protein crystals were used in X-ray diffraction experiments at the Diamond Light Source synchrotron facility (Oxford, UK). Data for SAD experimental phasing was collected at a wavelength of 1.28 Å and was processed using XDS. A single atom of Zn^2+^ (co-purified with BepA) was identified using SHELXD. This initial map was used for auto-building with Phenix. Models were improved by iterations of refinement using Phenix and manual manipulations in COOT.

### Consurf analysis

The consurf server was used to analyse conservation of surface residues. A multiple alignment of BepA homologues was generated using Clustal Omega and submitted to the consurf server along with the BepA pdb file as a basis for conservation analysis.

### Mutagenesis of bepA

Mutations in *bepA* were created using a PCR-based site directed mutagenesis approach utilising the pET20b::*bepA::6xHis* vector as template. Briefly, pET20b::*bepA::6xHis* vector was used in 18 cycles of PCR utilising the Phusion polymerase (NEB) as described by the manufacturer, but using complementary primers containing the desired mutation flanked by at least 15-bp of sequence. As a negative control, replica reactions were set up and the polymerase omitted. Template DNA was then digested by addition of 20 units (1 μl) *DpnI* restriction enzyme (NEB: R0176S) and incubation at 37°C for 1 h. The reaction mixture was then used to directly transform NEB DH5-α high-efficiency competent cells. Mutations were confirmed by plasmid isolation and Sanger sequencing (Source Bioscience).

### Functional screening of mutant bepA

Parent or *ΔbepA* cells were first transformed with the appropriate pET20b::*bepA::6xHis* vector and stored as glycerol stocks at −80°C. Starter cultures were generated by growth overnight (~16 h) at 37°C with aeration in LB broth (10 g/L tryptone; 5 g/L yeast extract; 5 g/L NaCl) supplemented with 100 μg/ml carbenicillin. Cells were normalised to OD_600_ = 1 and then 10-fold serially diluted before 1.5 μl was spotted onto the relevant LB agar plates. Cells were then incubated at 37°C overnight (~16 h) and the plates photographed for record. Cells were screened on LB agar plates supplemented with 100 μg/ml carbenicillin, vancomycin at the stated concentrations and 2 mM TCEP (tris(2-carboxyethyl)phosphine) where stated.

### Western blot

To examine the expression of BepA in *ΔbepA* or parent *E. coli*, cells were grown as described for the functional screening of mutant BepA experiments. The OD_600_ of the cultures was recorded and were isolated by centrifugation and resuspended in Laemmli buffer so that the number of cells in each sample was equivalent. Samples were boiled for 10 min before being resolved by SDS-PAGE and subjected to western blotting using anti-6xhis antibodies (TaKaRa: 631212) as primary antibody and HRP conjugated anti-rabbit (Sigma Aldrich: A6154) antibodies as secondary for detection by use of the ECL system.

### CPRG permeability assay

Following double transformation with the relevant pET20b::*bepA::6xHis* plasmid and the pRW50/CC-61.5 *lac* reporter plasmid, cells were grown to mid-exponential phase (OD_600_ 0.4-0.6) in LB broth with aeration and harvested by centrifugation. Cells were resuspended in LB broth to an OD_600_ of 0.1 and 5 μl used to inoculate 96-well culture plates containing 150 μl LB agar supplemented with CPRG (Chlorophenol red-β-D-galactopyranoside – Sigma) (20 μg/ml), carbenicillin (100 μg/ml) and tetracycline (15 μg/ml). 96-well plates were incubated at 30°C and the optical density 300-800 nm monitored every 20 min for 48 h. By using the absorbance of LacZ-strains unable to turn over CPRG, we created an estimating function that predicts the expected absorbance due to cell growth at 575 nm (CPR peak absorbance) using the absorbance at 450nm and 650nm. By subtracting the actual absorbance at 575 nm, from the expected growth-related absorbance we derive the CPRG turnover score, which is exclusive to cell membrane permeability. For both expected and measured absorbance at 575 nm, the timepoint of 24 h post-inoculation was used.

### LPS labelling, Lipid A isolation and analysis

Labelling of LPS, Lipid A purification, TLC analysis and quantification were done exactly as described previously (Chong et al., 2015). Briefly, starter cultures were incubated at 37°C overnight with aeration in LB broth supplemented with 100 μg/ml carbenicillin. Starter cultures were then subcultured into 5 ml LB broth supplemented with 100 μg/ml carbenicillin and the experiment completed precisely as described previously including the addition of the positive control, in which the parent strain was exposed to 25 mM EDTA for 10 min prior to harvest of cells by centrifugation in order to induce PagP mediated palmitoylation of Lipid A (Chong et al., 2015). Experiments were completed in triplicate and the data generated was analysed as described previously.

## Acknowledgements

We thank Professor Jeff Cole for stimulating discussion and assistance with manuscript editing. This work was funded by BBSRC and MRC grants to IRH. The lipid A palmitoylation work was supported by the Singapore Ministry of Education Academic Research Fund Tier 2 (MOE2013-T2-1-148) grant (to SSC).

## Author contributions

BepA was identified as a target for study by FM and the project was facilitated by IRH. The BepA mutant was made by FM. Production of the protein was completed by YS, whereas further production, purification, crystallisation and modelling of the BepA structure was done by ITC and ALL. Consurf analysis, mutagenesis and functional analysis screens were done by JAB with guidance from ALL. Lipid A palmitoylation assays were done by JAB and ZSC under the supervision of SSC. CPRG permeability assays were also completed by JAB with the advice and supervision of MB. CPRG assay data was processed and analysed by MB and GK. Manuscript preparation was completed by JAB and general project design was done by JAB with the guidance of ALL and IRH.

---

**Figure S1 - Comparison of PDB: 3C37 and BepA shows the active site lid**

Comparison of the *Geobacter sulfureducens* m48 protease structure PDB: 3C37 with that of the m48 protease domain of BepA presented here shows the active site lid formed by the missing residues H145-S195. **A.** The BepA structure is shown with the TPR domain in blue and the protease domain in white with the structure made transparent to facilitate visualisation of the 3C37 structure in red. **B.** Zoom in of the protease domain overlays from the BepA structure and that of 3C37 with important residues and structures labelled.

**Figure S2 - Western blot analysis of BepA::6xHis expression**

Analysis of BepA expression by western blot analysis. Cells carrying pET20b encoding WT or mutated copies of BepA in the parent or Δ*bepA* strain background were harvested and resuspended in Laemmli buffer so that the number of cells in each sample was equal. Proteins were separated by SDS-PAGE and transferred by western blot. The empty vector control is labelled EV. Western blotting was completed using anti-6xHis antibody raised in mice and anti-mouse::HRP secondary antibody to target the BepA::6xHis protein in samples used for vancomycin sensitivity screens.

